# Switching to one or the other : Shorebirds behavioural flexibility in food transport mechanisms

**DOI:** 10.64898/2026.07.29.741647

**Authors:** Marine Pery, Maëlle Rivain, Glenn Le Floch, Guillaume Gelinaud, Ophélie Deffes, Alexandre Petry, Michel Baguette, Vincent Bels

## Abstract

Shorebirds provide an excellent model for investigating the relationship between bill morphology and food acquisition. Food acquisition comprises three successive behavioural stages: foraging (locomotion and prey capture), feeding (food handling and transport), and swallowing. During food transport, these birds use two non-lingual mechanisms, surface-tension transport (ST) and ballistic transport (BT), whose characteristics depends on the kinematics of head and beak movements and the physical properties of the food. We investigated the behavioural flexibility of these transport mechanisms in captive and free-ranging individuals of two species with contrasting beak morphologies: the Pied avocet (*Recurvirostra avosetta*) and the Black-winged stilt (*Himantopus himantopus*). Although these species have beaks of a similar size, avocets are distinguished by their upward-curved beaks, whereas stilts have a rather straight beak. We examined the effects of food water content and the presence or absence of water in the beak on food transport by quantifying maximum gape, maximum head displacement, and maximum head velocity.

Transport kinematics were jointly influenced by food properties and water availability in the beak. Moist food improved transport performance in both species, whereas dry food required compensatory increases in gape amplitude and head movements, demonstrating that neither ST nor BT constitutes a fixed behavioural sequence. Species also differed consistently in their transport strategies: Black-winged stilts relied on slower, larger-amplitude head movements, whereas Pied Avocets exhibited faster, more precise movements, particularly when water was present.

These findings demonstrate that ST and BT share common biomechanical foundations while being governed by rapid kinematic adjustments to changing environmental conditions that probably correspond to flexible motor control. This behavioural flexibility in food transport is therefore likely to enhance feeding performance and ecological resilience in the heterogeneous habitats occupied by shorebirds, suggesting that context-dependent modulation of transport behaviour represents an important adaptive feature of these shorebirds.

## Introduction

Beak diversification has played a central role in avian evolution by enabling access to a wide range of trophic resources and ecological niches (1,2). Because the avian beak is directly involved in prey acquisition and processing, variation in beak morphology is closely associated with feeding ecology and functional specialization (3). Shorebirds (Charadriiformes) provide a particularly relevant model to investigate these relationships because they exploit highly diverse habitats and prey types using distinct feeding strategies (4–6). Across shorebirds, differences in beak size and shape are strongly associated with food-acquisition modes. Robust straight beaks are adapted for handling hard prey, elongated beaks facilitate probing for buried prey, and recurved beaks enable prey exploitation within the water column (7,8). These functional differences may also extend within species, as observed in oystercatchers where variation in beak shape is associated with alternative feeding specializations and prey preferences (9–12).

Food acquisition behaviour in shorebirds can be divided into three successive phases: foraging, feeding, and swallowing (4). Foraging includes prey search and capture, feeding corresponds to prey handling and transport from the beak tip to the pharynx, and swallowing represents the final ingestion phase. These phases involve distinct functional performances that are closely associated with morphology, particularly beak length and shape, following the performance-based framework (13–15).

The Recurvirostridae (avocets and stilts) provide a useful example of this functional integration. These species forage primarily in shallow aquatic habitats using continuous walking while capturing prey visually by pecking or tactilely by probing. Both groups also perform sweeping movements through water, although only avocets possess recurved beaks and lamellae allowing efficient filtration of suspended prey (16,17). Pied Avocets feed on fish, crustaceans, and aquatic insects (18,19), whereas Black-winged Stilts mainly consume aquatic invertebrates and insects (20).

Food transport in birds relies on coordinated gape modulation and head movements governed by stereotyped motor patterns and biomechanical constraints (21–24). Shorebirds use both lingual and non-lingual transport mechanisms to move prey from the tip of the beak toward the pharynx (4,25). Lingual transport depends on tongue movements maintaining continuous contact with the prey. In contrast, the two principal non-lingual transport mechanisms are surface tension transport (ST) and ballistic transport (BT). ST relies on water droplets enveloping the prey, with transport driven by capillary and surface-tension forces through coordinated mandibular movements and head oscillations (Fig 1) (22,23,26). In contrast, BT depends on rapid backward head movements accelerating the prey toward the pharynx while the prey loses contact with the beak (21,25). Because ST depends on stable droplets, small gape amplitudes, and capillary cohesion, its functional range may be strongly constrained by prey size, prey hydration, and environmental conditions (22).

**Fig 1.**
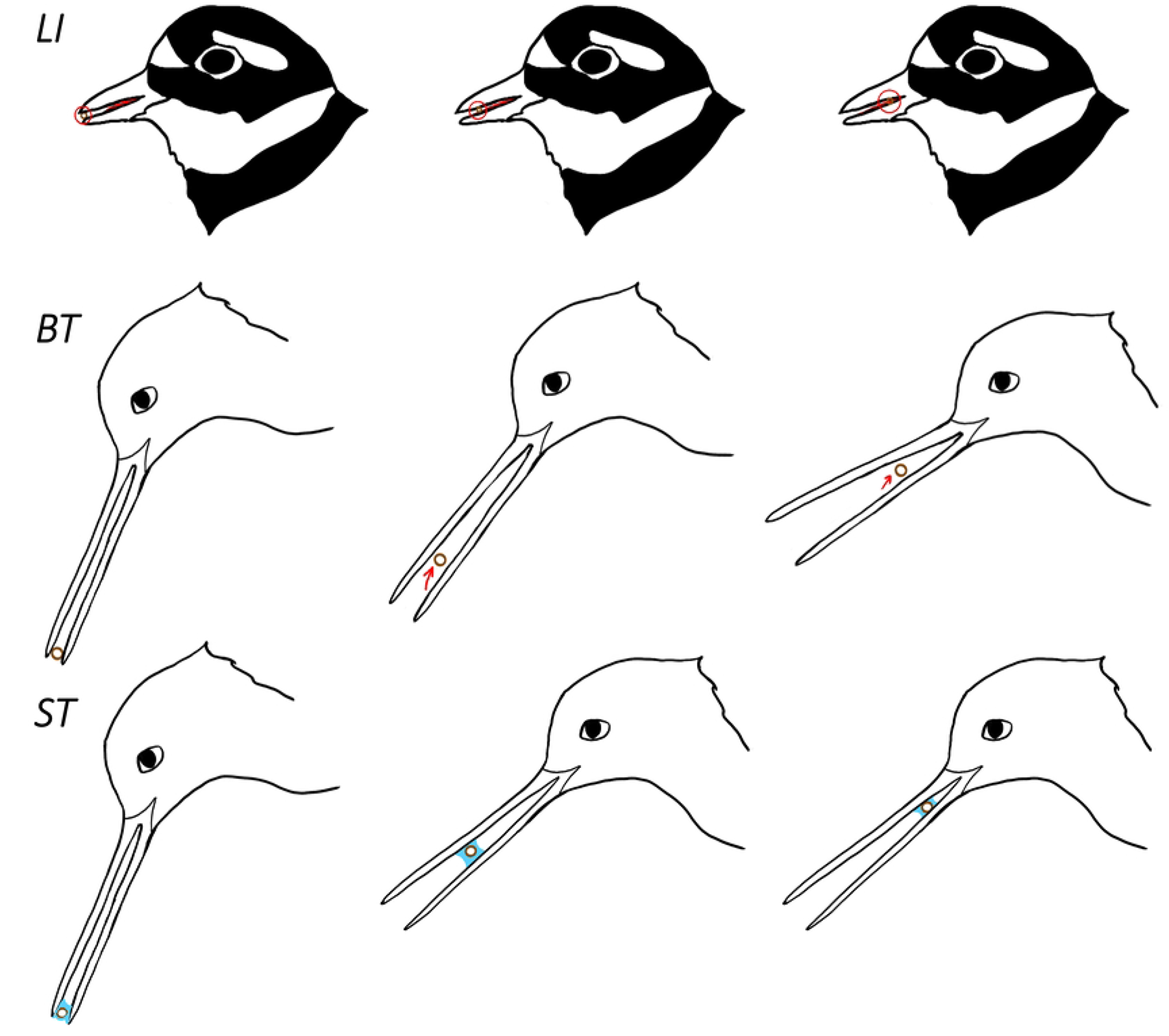
Schematic representation of food transport mechanisms. Food particle is presented as a brown circle, water with blue and tongue is red. LI lingual transport, BT ballistic transport and ST surface tension.

Understanding how shorebirds adjust transport behaviour to changing environmental conditions is particularly important because feeding performance may be affected by prey properties, water availability, and habitat alteration. Behavioural responses to prey characteristics have already been documented in shorebirds. For example, oystercatchers modify feeding behaviour according to prey size and profitability (27). Similarly, variation in food type, size, and consistency can directly affect feeding performance in birds (28,29). Such prey-dependent behavioural flexibility suggests that feeding performance may emerge from interactions between environmental conditions, prey properties, and biomechanical constraints.

Despite extensive work on shorebird feeding ecology, the kinematic basis of food transport remains poorly documented. To our knowledge, shorebirds are among the few avian groups in which distinct non-lingual transport mechanisms coexist within the same species and may be selected according to immediate environmental and prey-related contexts. Although ontogenetic and ecological changes in feeding behaviour have been described in other bird groups such as Anatidae (20,30,31), these transitions generally occur across developmental or ecological timescales rather than through rapid switching between distinct transport mechanisms during feeding. Food transport in shorebirds may therefore rely on context-dependent individual choice between transport modes rather than unrestricted variation within a single mechanism. Because ST and BT operate under different biomechanical and hydrodynamic constraints, environmental and prey-related contexts may determine which mechanism remains mechanically effective during feeding.

In this study, we investigate food transport kinematics during the feeding phase in two shorebird species of the family Recurvirostridae: the Black-winged Stilt (*Himantopus himantopus*) and the Pied Avocet (*Recurvirostra avosetta*). Both species have previously been shown to use ST and BT under natural conditions (4), making them suitable models to investigate transport-mode selection within a shared ecological context. We analyse how transport mechanisms vary according to water presence and prey hydration state. We predict that environmental and prey-related contexts modify the biomechanical effectiveness of each transport mechanism, thereby favouring context-dependent shifts between ST and BT.

## Materials and Methods

### Study species and data collection

Black-winged stilts (*Himantopus himantopus*) and Pied avocets (*Recurvirostra avosetta*) were observed and filmed both in captivity (*ex situ*) and natural conditions (*in situ*).

#### Ex situ

Black-winged stilts and Pied avocets are housed in a 2100m² aviary within the “Parc animalier et botanique de Branféré” (Le Guerno, France). The group is composed of five black-winged stilts (3 females, 2 males) and five pied avocets (1 female, 4 males). The aviary is also housing Eurasian spoonbills (*Platalea leucorodia*), Squacco herons (*Ardeola ralloides*), Little egrets (*Egretta* garzetta), Marbled ducks (*Marmaronetta angustirostris*), Oystercatchers (*Haematopus ostralegus*) and Ruffs (*Calidris pugnax*).

Food is composed of standard zoological bird food, namely microfloating (0.00060g, 2.27mm diameter), “regular” floating (0.084g, 6.82mm diameter), superworms (*Zophobas morio)* (0.30g, 4-6cm length), mealworms (*Tenebria molitor*) (0.1g, 1.20-1.80cm length) and “insect powder” (Nutribird, “insect pâtée”, about 0.05 g and 5-10 mm diameter), i.e., crushed insects, crustaceans and mollusks. Microfloating was always soaked in water whereas other food sources were dry. Superworms and mealworms were considered as treats and not always distributed. Birds can also hunt in a natural pond occupying 75% of the aviary, with water height from 15 cm to 1.4 m.

We collected 23 videos of individuals eating from a feeding bowl or from the pond, from which we analyzed 64 sequences (31 for Black-winged stilts, 33 for Pied avocets, supplementary material S1) where complete mandibular spreading cycles were seen. By complete mandibular spreading cycles we mean complete sequence of food transportation from the tip of the beak to the pharynx. Videos were recorded from 25 to 60 fps with a Lumix DC-GH5, lens Leica DG Vario-Elmar 100-400mm and lasted minimum 1 minute.

#### In situ

Black-winged stilts were mostly filmed in august 2022, in the « Domaine des oiseaux » Nature Reserve, Mazères, Ariège, France. Pied avocets were filmed in different locations : the « Teich » Nature Reserve, Gironde, France; Salin-de-Giraud, Bouches-du-Rhône, France, the « Marais de Séné » Nature Reserve, Morbihan, France and the « Marais de Suscinio » Morbihan, France in 2024 and 2025. Videos were recorded at 30 or 60 fps with an Olympus OMD EM1x with lens Zuiko Pro 150-400mm and lasted minimum 1 minute.

Food within the beak was sometimes difficult to correctly identify. Therefore we noted food categories as worms, *Artemia* sp. or small invertebrates (i.e., all small invertebrates that could not be discriminated on video).

In total for both species, 31 videos were selected for analyses from which 58 sequences (30 for black-winged stilts, 28 for pied avocets) showing complete mandibular spreading cycles were extracted. We considered every different video as a new individual except when several birds could be seen and distinguished on one video.

### Video analysis

First, we performed a visual description of the food acquisition behaviours. We assigned on each video the different phases of the foraging and feeding behaviours. We used the repertoire of food acquisition behaviours in shorebirds (4), in which foraging describe the way birds move (locomotion) and catch food particles (capture), whereas feeding describe the way birds handle (handling) and move food particles from the tip of the beak to the pharynx (transport). Second, we focused on the transport phases identified at the previous step. We analyzed food particles trajectories using Kinovea (https://www.kinovea.org/, version 2023.1.2). For both conditions (*ex situ* and *in situ*), sequences started at the first image before any food particle started to be transported to the pharynx and finished when no more visible (i.e. swallowed or caught by the tongue before swallowing). Sequences were chosen so that the bird was seen from its profile, in order to follow the trajectory of the particle precisely.

Using Kinovea, five points were placed on the first image of the videos : two on the beak (superior extremity and inferior extremity), one on the food particle, one on the eye of the bird and one at the back of its head. To standardize the measurements we used the mean for beak length measured as a straight line from the culmen to the tip of the beak. Measurements were made on 6 specimens of each species in the collection of National Museum of Natural History, France. Those values (mean=6.27 cm, sd=0.43 for *H. himantopus* and mean=8.42 cm, sd=0.66 for *R. avosetta*) were used for scaling the trajectories of food particles in the videos. Frame of reference was calibrated as aligned with the image axis (**see S?**). The position of the different points throughout the sequence was checked in order to prevent any error in measurements. Then, x and y’s position values of each five points were extracted from Kinovea. Gape (*i.e.* distance between superior and inferior mandible) was calculated using *G* 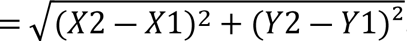.

X1-Y1 and X2-Y2 being the x and y coordinates of the beak superior extremity and inferior extremity respectively. Using the coordinates from the eye, we determined head movement (in cm) using the same formula, comparing the coordinates between the first frame and the next. Using time (ms) and the distance calculated in head movement (cm), we determined the head speed at which the head was moving from one frame to another (cm/ms).

Maximum gape (MG), maximum head movement (MHM) and maximum head speed (MHS) were measured for each sequence and extracted for statistical analyses. Type of transport, either surface tension (ST) or ballistic transport (BT) for the sequence was also noted, as well as, presence or absence of water in the beak during the sequence. As for food hydric content, we considered insect powder plus floating, micro-organisms plus worms and microfloating plus artemia as dry, moist and wet food, respectively.

### Statistical analyses

Using RStudio (version 2025.05.1+513), maximal gape (MG), maximum head movement (MHM) and maximum head speed (MHS) were analyzed for each sequence (see **S**? for data).

Firstly, we tested for data normality. As their distribution was not normal, we used non-parametric statistics. One outlier was removed for analysis, identified as an error of measurement in Kinovea. A total of 121 sequences were extracted for statistical analyses.

Then, we investigated intraspecific and interspecific differences in MG, MHM and MHS. A Mann Whitney-U (MWU) test was used to investigate for differences in the tested variables regarding the presence or absence of water droplets in the beak during transport. As for food hydric content we investigated the differences using a Kruskal-Wallis test (‘stats’ package v. 4.4.1). Post-hoc Dunn test using a Bonferonni correction was then computed to look for interactions between food hydric content categories (i.e., dry moist and wet). MG, MHM and MHS were also tested for differences between transport type (i.e., BT or ST) using an MWU.

Interspecific differences were studied using a Mann Whitney-U test to compare *H. himantopus* and *R. avosetta* MG, MHM and MHS when water droplet was present in the beak and not. Another MWU test was computed to compare the three variables when species were eating either dry, moist or wet food. Lastly, differences between species was tested in BT and ST transport using a MWU test. Graphs were generated using RStudio and the ggplot2 package (v 4.0.0).

## Results

We observed two different types of BT. When the food particle was too large to only use ST but the presence of water allowed its motion induced by a form of capillarity, birds use a form of BT using both water and inertia, corresponding to the straight-draw transport described by (4). This transport mechanism is represented by sequences where water droplet in the beak is combined with BT (Table 1).

**Table 1.** Number of sequences regarding species, water droplet presence, food hydric component and food transport mechanism.

| Transport mechanism | BT | ST | Total |
| --- | --- | --- | --- |
| **Black-winged stilt** | 30 | 31 | 61 |
| **No water in the beak** | 21 | 0 | 21 |
| Dry food | 17 | 0 | 17 |
| Moist food | 4 | 0 | 4 |
| **Water in the beak** | 9 | 31 | 40 |
| Moist food | 8 | 26 | 34 |
| Wet food | 1 | 5 | 6 |
| **Pied avocet** | 13 | 47 | 60 |
| **No water in the beak** | 8 | 0 | 8 |
| Dry food | 4 | 0 | 4 |
| Moist food | 4 | 0 | 4 |
| **Water in the beak** | 5 | 47 | 52 |
| Moist food | 5 | 18 | 23 |
| Wet food | 0 | 29 | 29 |
| **Total** | 43 | 78 | 121 |

BT : ballistic transport; ST : surface tension.

We determined the maximum gape (MG) measurements for both species using either BT or ST transport mode (Fig 2).

**Fig 2.**
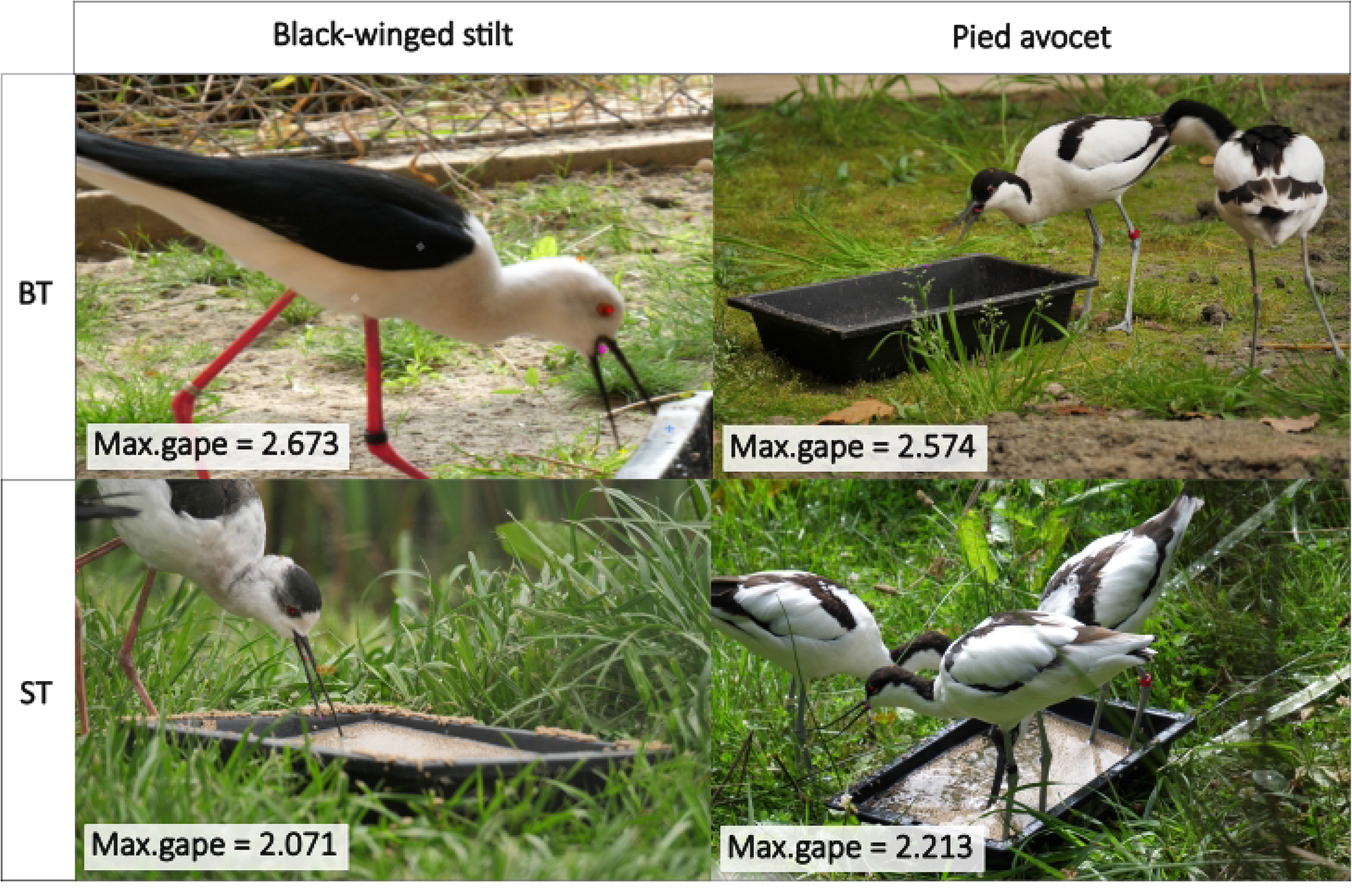
Maximum gape (MG, cm) measured for Black-winged stilts and Pied avocets for sequences with BT and ST transport mechanisms.

## Intraspecific comparisons

### Maximum gape

When water droplets were not observed within the beak during food transport, MG was significantly higher than with water droplets for Black-winged stilts and Pied avocets (MG*water availability, p<0.001 and p<0.01, respectively for each species, Fig 3-A). MG was significantly different regarding food hydric content for Black-winged stilts and Pied avocets (MG*food hydric content, p<0.01 and p<0.001, respectively for each species, Fig 3-B). When eating dry food, maximum gape of Black-winged stilts was significantly higher than for moist food (post-hoc, MG*food(dry-moist), p<0.001, Fig 3-B). Meanwhile MG was significantly higher when eating wet food compared to moist food (post-hoc, MG*food(moist-wet), p<0.01, Fig 3-B). There was no significant difference of maximum gape regarding food hydric content for Pied avocets (MG*food hydric content, p=0.214).

**Fig 3.**
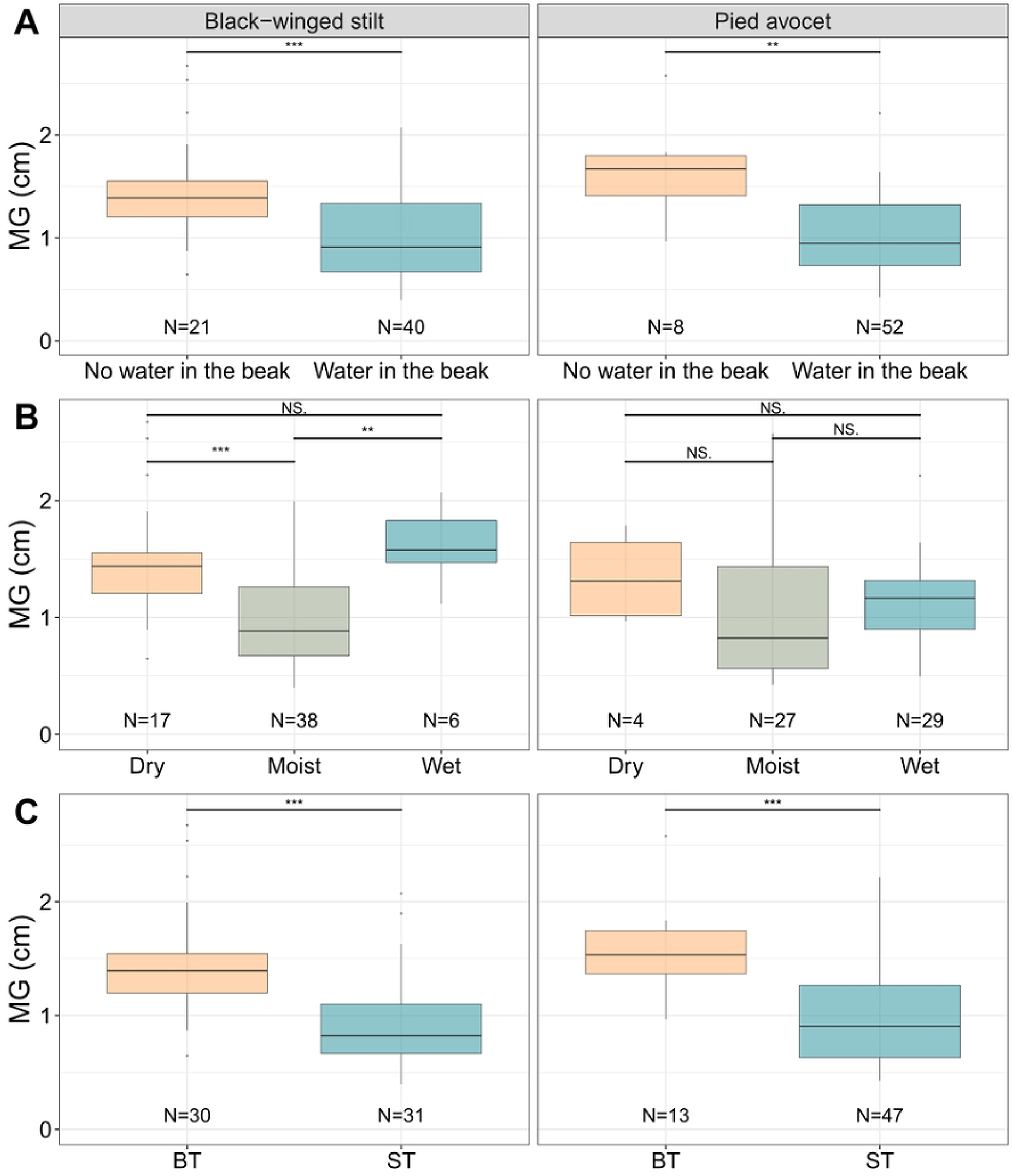
Maximum gape (MG, cm) measured for Black-winged stilts and Pied avocets. (A) MG comparison when water was present or not in the beak during food transport. (B) MG differences regarding humidity of the food source (dry, moist or wet). (C) is MG compared between BT and ST transport mechanisms. (A&C : Mann-Whitney tests; (B) : Kruskall-Wallis tests and Dunn tests for post-hoc comparisons; NS : non-significant, * : p<0.05, ** : p<0.01, *** : p<0.001).

Black-winged stilts and Pied avocets had a significantly higher MG using ballistic transport (BT) mechanism than when using surface tension (MG*transport, p<0.001 and p<0.001 respectively for Black-winged stilts and Pied avocets, Fig 3-C). Using surface-tension mechanism, both species had a significantly higher MG while eating wet food (MG*food(moist-wet) with ST, p<0.001 and p<0.001).

### Head movement and speed

For Black-winged stilts, water presence within the beak and transport type significantly impacted MHM but not MHS. MHM was significantly higher with water in the beak than without (MHM*water availability, p<0.01, Fig 4-A). MHS was not significantly different with or without water in the beak (MHS*water availability, p=0.522, Fig 5-A).

**Fig 4.**
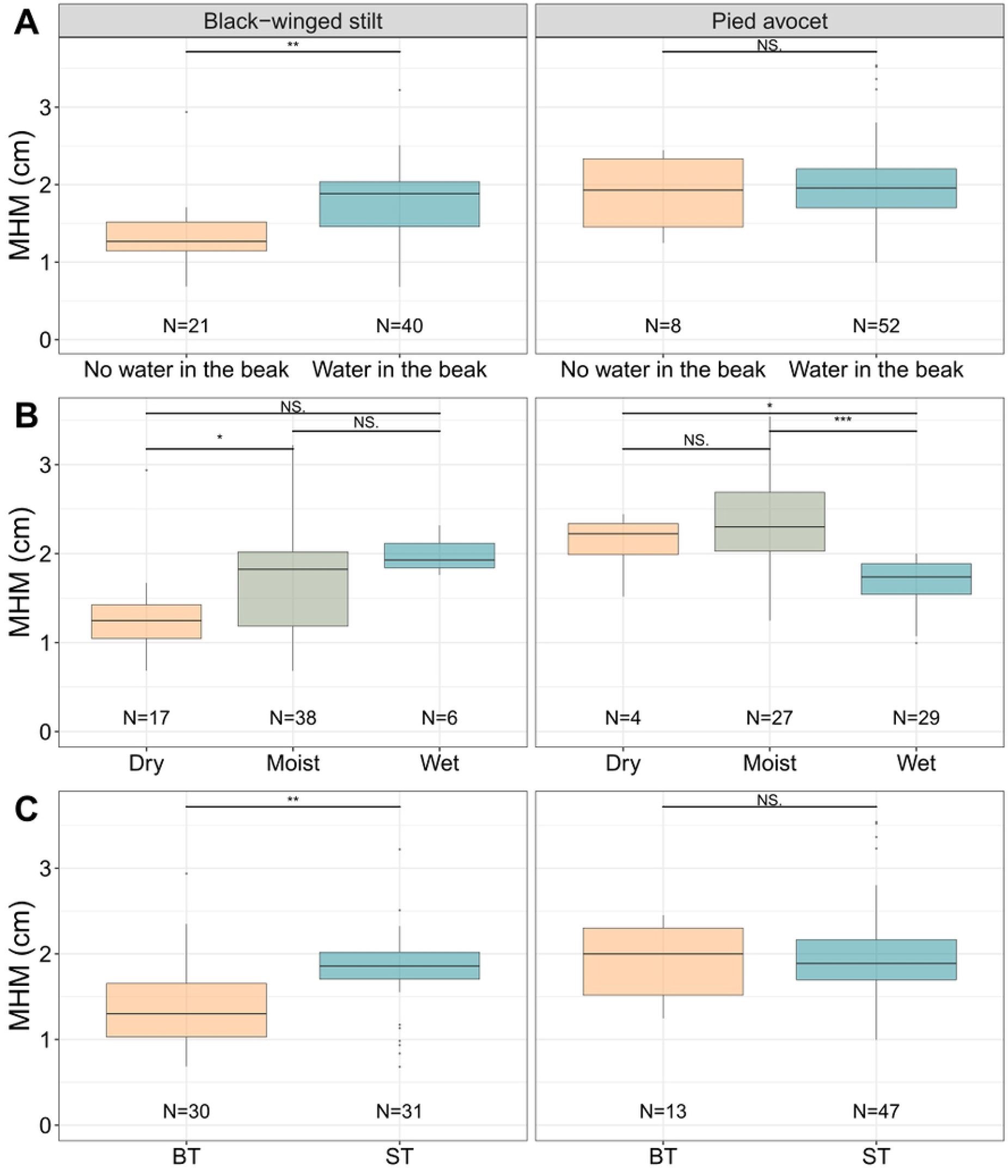
Maximum head movement (MHM, cm) measured for Black-winged stilts and Pied avocets. (A) MHM comparison when water droplet is present or absent from the beak. (B) MHM differences regarding humidity of the food source (dry, moist or wet). (C) is MHM compared between BT and ST transport mechanisms. (A&C : Mann-Whitney tests; (B) : Kruskall-Wallis tests and Dunn tests for post-hoc comparisons; NS : non-significant, * : p<0.05, ** : p<0.01, *** : p<0.001).

**Fig 5.**
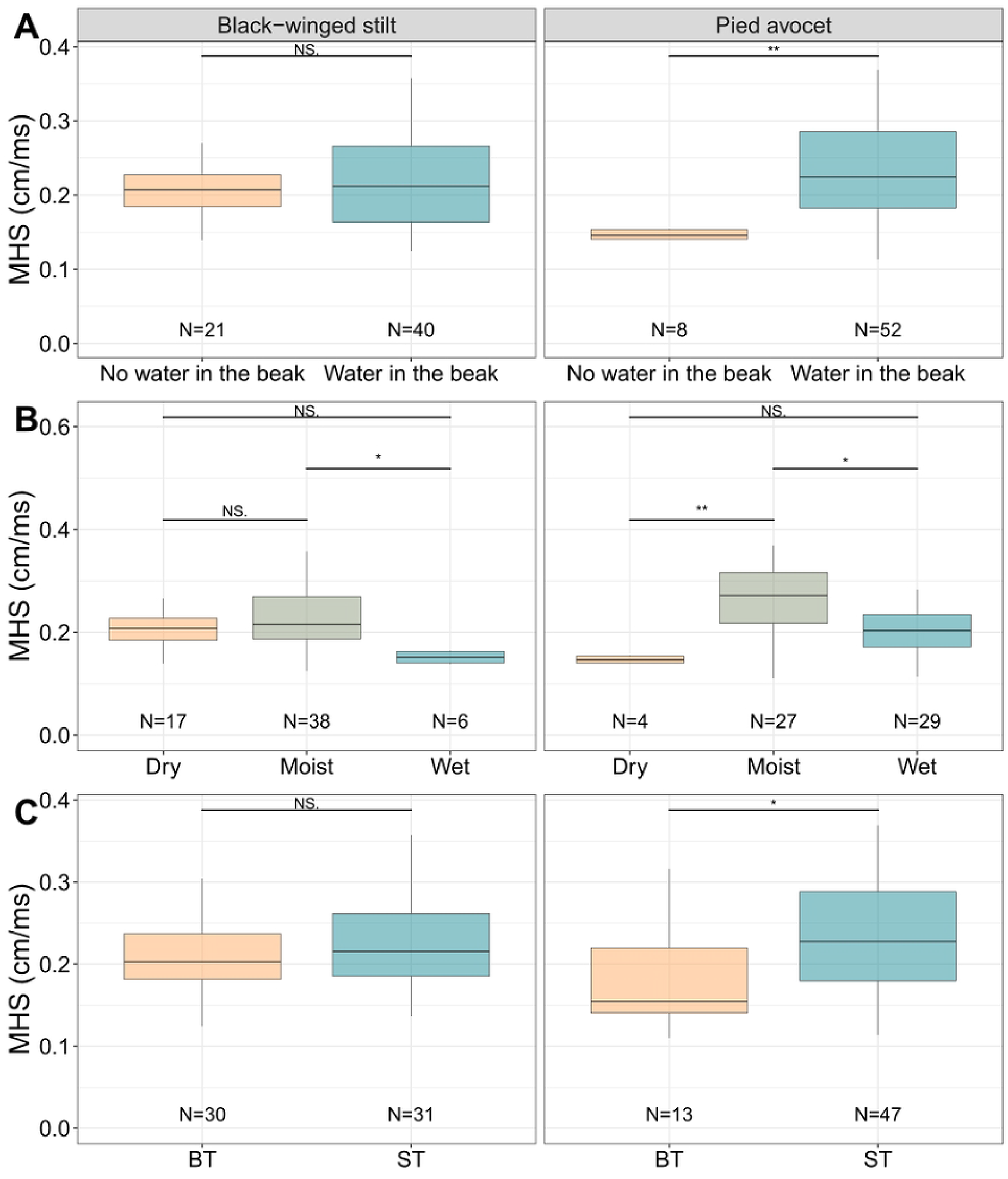
Maximum head speed (MHS, cm/ms) measured for Black-winged stilts and Pied avocets. (A) MHS comparison when water droplet is present or not in the beak. (B) MHS differences regarding humidity of the food source (dry, moist or wet). (C) is MHS compared between BT and ST transport mechanisms. (A&C : Mann-Whitney tests; (B) : Kruskall-Wallis tests and Dunn tests for post-hoc comparisons; NS : non-significant, * : p<0.05, ** : p<0.01, *** : p<0.001).

Results are reversed for Pied avocets, MHS was significantly impacted but not MHM. MHS was significantly higher when water was in the beak than not (MHS*water availability, p<0.01, Fig 5-A). MHM was not significantly different with or without water in the beak (MHM*water availability, p=0.704).

Food hydric content had an impact on MHM and MHS for both species (MHM*food hydric content, p<0.01 and p<0.001; MHS*food hydric content, p<0.05 and p<0.001, respectively for Black-winged stilts and Pied avocets).

When eating moist food, MHM was significantly higher than when eating dry food for Black-winged stilts (MHM*food(dry-moist), p<0.05, Fig 4-B). Meanwhile, MHS was significantly lower when eating wet food than when eating moist food (MHS*moist-wet, p<0.05, Fig 5-B).

For Pied avocets, MHM was significantly lower when eating wet food than when eating dry or moist food (MHM*wet-dry, p<0.05; MHM*wet-moist, p<0.001, Fig 4-B). In the meantime, MHS was significantly higher when eating moist food than when eating dry or wet food (MHS*moist-dry, p<0.01; MHS*moist-wet, p<0.05, Fig 5-B).

Regarding transport use by Black-winged stilts, MHM was significantly higher for surface tension compared to ballistic transport (MHM*transport, p<0.01, Fig 4-C) but MHS was not significantly different between transport mechanisms (MHS*transport, p=0.236, Fig 5-C). For Pied avocets MHS was higher for surface tension transport than ballistic transport (MHS*transport, p<0.05, Fig 5-C). MHM was not significantly different for transport mechanisms in this species (MHM*transport, p=1, Fig 4-C).

Using surface-tension mechanism, Black-winged stilts did not show significant differences in MHM comparing food hydric content but had a significantly higher MHS while eating moist food (MHM*food(moist-wet) with ST, p=0.658; MHS*food(moist-wet) with ST, p<0.05).

For Pied avocets MHM and MHS were significantly higher while eating moist food (MHM*food(moist-wet) with ST, p<0.001; MHS*food(moist-wet) with ST, p<0.001).

Using ballistic transport, MHS was significantly higher when water was present in the beak for Pied avocets but not for Black-winged stilts (MHS*water availability, p<0.05).

## Interspecific comparisons

### Maximum gape

There was no difference of MG between species when comparing water presence in the beak (MG*species(water), p=0.324 and p=0.940 for without and with water respectively).

When eating wet food, maximum gape was higher for Black-winged stilts compared to Pied avocets (MG*species(wet food), p<0.01). No other significant difference was noted (MG*species(dry food), p=0.763; MG*species(moist food), p=0.984).

Regarding transport, there was no difference of maximum gape between species in BT and ST food transport (MG*species(transport), p=0.289 and p=0.522 for BT and ST respectively).

### Head movement and speed

MHM and MHS were significantly different between Black-winged stilts and Pied avocets when water was not in the beak. MHM was significantly higher for Pied avocets without water in the beak, meanwhile MHS was significantly higher for Black-winged stilts (MHM*species(no water in the beak), p<0.05 Fig 7-A; MHS*species(no water in the beak), p<0.01, Fig 8-A). There was no significant difference of MHM and MHS between species when water was in the beak (MHM*species(water in the beak), p=0.165 Fig 7-A; MHS*species(water in the beak), p=0.165 Fig 8-A).

MHM of Pied avocets was significantly higher than Black-winged stilts when eating dry and moist food, but the opposite when eating wet food (MHM*species(dry food), p<0.05; MHM*species(moist food), p<0.001; MHM*species(wet food), p<0.05, Fig 7-C).

MHS of Pied avocets was significantly higher than Black-winged stilts when eating moist and wet food, but the opposite when eating dry food (MHS*species(moist food), p<0.05; MHS*species(wet food), p<0.05; MHS*species(dry food), p<0.05, Fig 8-C).

When comparing ballistic transport between species, MHM of Pied avocets was significantly higher than Black-winged stilts (MHM*species(BT), p<0.01, Fig 7-B). There was no significant difference of MHM between species for ST (MHM*species(ST), p=0.329, Fig. 7-B) nor of MHS when comparing BT and ST transport mechanisms (MHS*species(BT), p=0.339; MHS*species(ST), p=0.290, Fig 8-B).

Using BT, MHM was significantly higher for Pied avocets when water was absent from the beak (MHM*species(no water in the beak) with BT, p<0.05). MHM was also significantly higher for Pied avocets when eating dry and moist food (MHM*species(dry food) with BT, p<0.01; MHM*species(moist food) with BT, p<0.001).

MHS was significantly higher for Black-winged stilts when water was absent from the beak (MHS*species(no water in the beak) with BT, p<0.01).

## Discussion

Food transport in shorebirds is strongly shaped by interacting morphological, ecological, and biomechanical constraints. Comparative studies indicate that most species rely on only one or two transport mechanisms, suggesting that feeding performance is constrained by both morphology and the physical requirements of food transport. (4) demonstrated that, across Western Palearctic shorebirds, each species employs at most two transport modes, often as part of a broader behavioural syndrome linking locomotion, prey capture, and food transport. This finding indicates that transport behaviours are integrated within coordinated functional systems rather than representing independent behavioural traits. Similarly, the ecomorphological framework proposed by (7) and formalized by (5) highlights strong associations between beak morphology and foraging strategy. Because food transport occurs downstream of prey capture, transport mechanisms are indirectly constrained by morphological specialization. Species with long, slender beaks adapted for probing, for example, possess anatomical features facilitating lingual intra-beak transport, thereby reducing dependence on alternative transport strategies. Biomechanical studies further indicate that transport mechanisms operate under strict physical constraints. (22) demonstrated that surface-tension transport (ST) depends on highly specific hydrodynamic conditions involving stable millimetric droplets, small gape amplitudes, and the transport of very small prey suspended within water droplets. Consequently, ST appears restricted to prey types and feeding conditions compatible with capillary transport. In contrast, ballistic transport (BT) relies on inertial acceleration generated by rapid cranial movements and is less dependent on water-mediated adhesion. Taken together, these findings suggest that transport flexibility in shorebirds may not primarily arise from unrestricted kinematic plasticity within a single mechanism, but rather from switching between distinct transport modes that remain individually constrained by biomechanics.

The comparative analysis presented here provides an opportunity to assess how the presence of water droplets and the moisture content of food influence transport kinematics and transport-mode expression in our study species. Our results show that food transport is governed by cranial kinematics (i.e., maximum gape, maximum head movement, and movement speed), prey biomechanical properties, and the presence of water within the beak. However, rather than indicating unrestricted behavioural modulation, our findings suggest that individuals select between two mechanically constrained transport mechanisms whose effectiveness depends on environmental and prey-related conditions. Both species reduce maximum gape when water droplets are present within the beak, suggesting that water facilitates food transport (Fig 3). In capillary feeding systems, prey items are transported while suspended within water droplets, indicating that droplet stability itself constitutes a critical component of transport efficiency (22). Surface-tension transport therefore requires small opening angles to maintain droplet cohesion and prevent droplet breakup. The reduced gape observed in our study is fully consistent with these physical constraints. This supports the hypothesis that surface tension and water capillarity assist in moving food items toward the pharynx while reducing the need for large mandibular excursions. Similar mechanisms have been described in *Phalaropus lobatus*, which exploits surface tension to transport food within water columns (23). Our findings therefore suggest that the presence of water does not merely alter transport kinematics but instead favours the expression of a distinct transport mechanism operating under specific biomechanical constraints (Figs 3, 4 and 5).

Food moisture content also strongly affects transport behaviour. Moist food was consistently associated with enhanced transport performance across several kinematic variables, whereas dry food required compensatory movements to maintain prey control. Wet food, in contrast, may generate instability associated with droplet cohesion and hydrodynamic transport dynamics (Figs 3, 4 and 5). These results suggest that individuals operate within biomechanical limits defining the conditions under which each transport mechanism remains mechanically effective. Rather than reflecting unrestricted variability within transport modes, differences observed here likely indicate context-dependent switching between ST and BT according to prey moisture, prey physical properties, and water availability. The variability observed here therefore expands upon earlier mechanistic studies by showing that birds exploit distinct transport mechanisms in response to changing ecological conditions (24).

Species-specific differences further support the existence of distinct transport strategies. The presence of water droplets appears to influence Black-winged Stilts toward slower movements with larger amplitudes (Figs 6 and 7), whereas Pied Avocets perform faster and more precise transport movements under similar conditions. This pattern may reflect a speed–accuracy trade-off similar to that described in granivorous songbirds, in which individuals optimize movement speed rather than maximize it to maintain feeding accuracy (28). Pied Avocets typically feed by sweeping their recurved beaks laterally through water, whereas Black-winged Stilts primarily peck and probe using straight slender beaks (16,17,32,33). These morphological differences likely constrain force application within the beak and may therefore influence not only transport kinematics but also the conditions under which one transport mechanism becomes more effective than the other. The association between beak morphology, biomechanical performance, and ecological function supports the view that avian beaks represent highly integrated feeding structures shaped by both ecological specialization and mechanical constraints (2,5,34).

**Fig 6.**
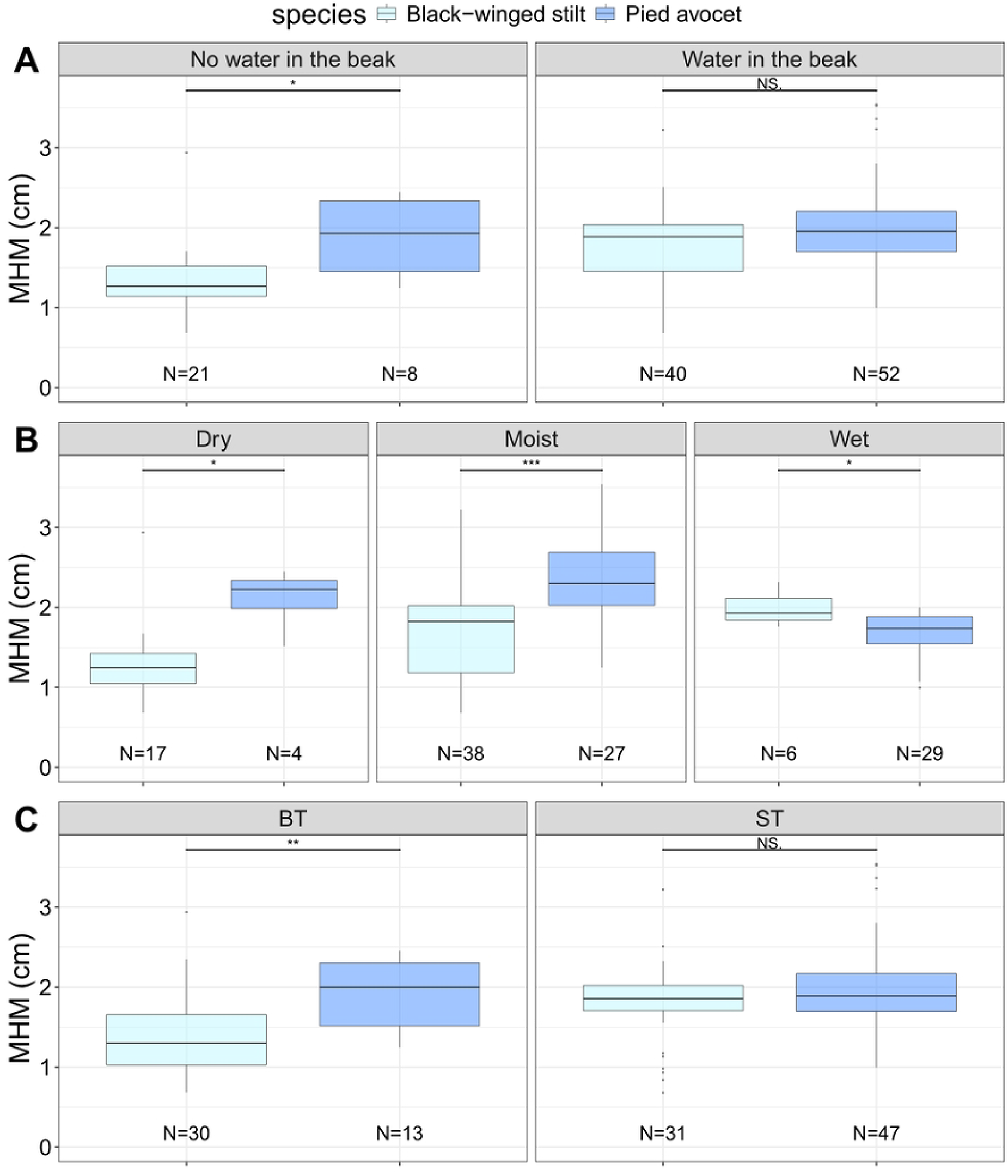
Maximum head movement (MHM, cm) comparisons between species, Black-winged stilts and Pied avocets. (A) MHM comparison between species when water was present or not in the beak during food transport. (B) MHM difference between species when eating dry, moist or wet food source. (C) MHM comparison between species when using BT or ST transport mechanism. (A&C : Mann-Whitney tests; (B) : Kruskall-Wallis tests and Dunn tests for post-hoc comparisons; NS : non-significant, * : p<0.05, ** : p<0.01, *** : p<0.001).

**Fig 7.**
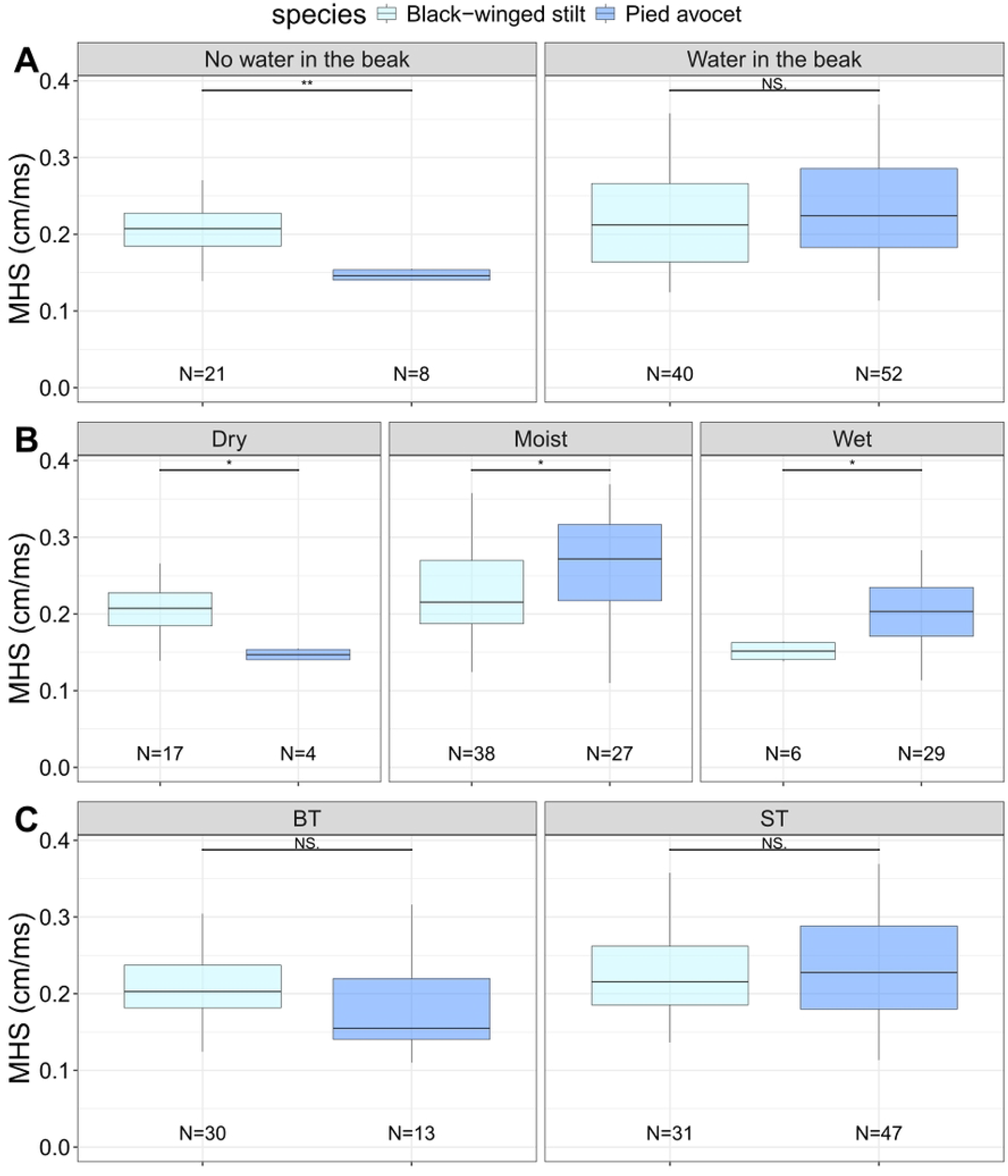
Maximum head speed (MHS, cm/ms) comparisons between species, Black-winged stilts and Pied avocets. (A) MHS comparison between species when water was present or not in the beak during food transport. (B) MHS difference between species when eating dry, moist or wet food source. (C) MHS comparison between species when using BT or ST transport mechanism. (A&C : Mann-Whitney tests; (B) : Kruskall-Wallis tests and Dunn tests for post-hoc comparisons; NS : non-significant, * : p<0.05, ** : p<0.01, *** : p<0.001).

The contrast between BT and ST therefore highlights behavioural switching between transport modes rather than unrestricted modulation within a single transport mechanism. BT is characterized by larger gape amplitudes and reduced dependence on water, providing a mechanically robust transport strategy when capillary transport conditions are no longer satisfied. In BT, the stereotypy of cranial and beak movements produces stable and repeatable cycles, emphasizing the invariant biomechanical properties of this mechanism (21,24,35). In contrast, ST depends on highly constrained hydrodynamic conditions involving stable droplets, small gape amplitudes, and water-mediated adhesion (22). Consequently, both transport mechanisms appear to function as relatively constrained motor modules rather than continuously variable kinematic systems (Figs 3, 4 and 5). The transition from ST to BT may therefore occur when the physical limits required for capillary transport are exceeded, particularly when droplet cohesion becomes unstable, gape amplitude increases, or prey properties no longer permit efficient surface-tension transport, as predicted by the hydrodynamic constraints described by (22).

Such switching behaviour may result directly from the physical limitations of ST transport. When prey size, prey moisture, droplet stability, or gape amplitude exceed the biomechanical conditions necessary for capillary ratcheting, birds may instead rely on BT, which is less dependent on capillary stability and water-mediated adhesion. Ballistic transport may therefore represent a mechanically robust fallback strategy operating outside the functional range of capillary transport. Similar prey-dependent switching between feeding techniques has been described in oystercatchers, which modify feeding behaviour according to prey characteristics and environmental conditions, highlighting the adaptive value of behavioural flexibility in food acquisition (36). This capacity to switch between transport mechanisms may represent an important adaptation for exploiting heterogeneous wetland environments where prey properties and water availability fluctuate continuously across spatial and temporal scales.

This interaction between constrained biomechanics and behavioural switching aligns with integrative frameworks proposed for avian feeding systems (33). These frameworks suggest that feeding behaviour relies on evolutionarily conserved neuromotor modules whose expression depends on morphology and environmental context. Recent studies in Passeriformes similarly demonstrated that feeding performance results from interactions among kinematic constraints, motor control, and acquired feeding skills (28). Our results support this interpretation by suggesting that behavioural flexibility in shorebird food transport primarily emerges through differential activation of distinct motor modules rather than through unrestricted variation within individual transport mechanisms. Although behavioural and feeding transitions have also been described in other avian groups such as Anatidae, these changes generally occur across ontogenetic or ecological timescales and are associated with developmental modifications of feeding systems (20,30,31). In contrast, our results suggest that shorebirds may rapidly switch between mechanically distinct transport mechanisms within the same feeding context according to immediate environmental and prey-related conditions. (4), similarly showed that shorebirds share a common organization of locomotion, prey capture, and transport, while retaining species- and context-dependent differences in behavioural expression. We observed clear differences between stilts and avocets under similar environmental conditions, suggesting that species differ not only in transport kinematics but also in the conditions favouring one transport strategy over another. Black-winged Stilts tend to favour larger gape amplitudes and slower transport movements, whereas Pied Avocets perform faster and more precise movements. This duality may reflect differences in beak shape, curvature, and feeding ecology. Similar relationships between morphology, motor control, and feeding performance have also been described in other bird groups, including Passeriformes and granivorous birds (29,37).

Overall, the present study demonstrates that food transport in shorebirds emerges from an interaction between fixed biomechanical constraints and context-dependent behavioural switching. Surface-tension transport and ballistic transport should not be interpreted as interchangeable variants of a single transport system but rather as distinct and complementary transport mechanisms operating under different physical and ecological constraints. Surface-tension transport is highly efficient under conditions permitting droplet stability and capillary ratcheting, whereas ballistic transport provides a mechanically robust alternative when these conditions are no longer satisfied.

Shorebirds appear to dynamically select the transport strategy that remains mechanically effective under current feeding conditions. Such behavioural switching may represent an important adaptation for maintaining feeding performance in heterogeneous wetland environments characterized by fluctuating prey properties and environmental variability.

More broadly, these findings support the view that avian feeding systems combine conserved neuromotor organization with flexible behavioural expression through motor-module selection. This interaction between mechanical constraints and behavioural switching may help explain how species maintain feeding efficiency despite environmental variability and changing ecological conditions. The ability to switch between transport mechanisms may therefore represent an underappreciated component of ecological resilience in shorebirds.

Future studies should investigate the sensory and biomechanical mechanisms governing transitions between transport modes, particularly the physical thresholds at which ST becomes unstable and BT is favoured instead. Experimental manipulations of prey size, prey surface properties, water availability, and beak kinematics would help clarify how environmental and morphological constraints interact to determine transport-mode selection. Comparative analyses across additional shorebird species may also reveal whether transport switching represents a generalized evolutionary strategy associated with ecological generalism or a derived adaptation restricted to specific feeding ecologies.

## Acknowlegments

We thank all the staff from the Parc animalier et botanique de Branféré (Le Guerno, France), Aurélie, Gwladys, Alice, Paul, who provided help when needed to film the animals (SJ 1219-24). We particularly thank Eleanor Lacassagne for her master thesis work in the MNHN collections. We also thank the Fondation de France for funding us (SJ 9922-24) and all collaborators from Bretagne Vivante-SEPNB (SJ 454-25). Michel Baguette is member of the Laboratory of Excellence TULIP funded by the ANR under grant ANR10-LABX-41.

## Supporting information

**S1 Fig. Frame of reference in a Black-winged stilt sequence.** Frame is aligned with the culmen (red lines). White line is the standardized beak measure used for calibration. Orange, red, purple, blue and yellow points are respectively following movement from the head, eye, food particle, superior and inferior beak.

**S2 Table. Sequences dataset**. Xlxs file presenting sequence number, species, transport, food hydric component, water presence, time start and end of the sequence, MG, MHM and MHS.

**S3 Table. Statistics dataset**. Xlxs file presenting variables tested, method, p value associated, statistics associated, and cross-comparisons with statistics, p values unadjusted and adjusted.

